# Post-assembly proteostasis of the respiratory chain is independent of respirasome organization

**DOI:** 10.64898/2026.10.13.758515

**Authors:** Jelena Misic, Pablo Rivera-Mejías, Clara Ursula Neumann, Nico Hüttmann, Pauline Michon, Dusanka Milenkovic, Ilian Atanassov, Xinping Li, Mikhail M. Savitski, Nils-Göran Larsson

## Abstract

Respirasomes have long been proposed to stabilize their constituent complexes. Yet the close relationship between individual complex biogenesis and respirasome formation has made it difficult to determine whether respirasomes are required for complex maintenance after assembly. Here, we uncoupled these processes using a genetic perturbation of the CI–CIII_2_ contact site that severely depletes respirasomes while preserving individual complex assembly and function. Across orthogonal measures of protein stability, turnover and solubility, respiratory-chain maintenance remained largely preserved despite severe respirasome depletion in mouse embryonic fibroblasts, even under mitochondrial stress. We further tested whether a requirement for respirasomes emerges *in vivo* when OXPHOS biogenesis is progressively impaired through tissue-specific TFAM deletion. Respirasome depletion did not accelerate the loss of CI, CIII_2_ or CIV during disease progression. Together, our findings show that respirasomes are not a major determinant of post-assembly respiratory-chain complex maintenance, even when mitochondrial homeostasis and OXPHOS biogenesis are challenged.

## Introduction

Respiratory chain complexes coexist as individual entities and higher order supercomplex assemblies in the inner mitochondrial membrane. Respirasomes comprise complex I (CI) and the dimeric complex III (CIII_2_) associated with one or more copies of complex IV (CIV) and can adopt more complex architectures, as revealed by *in situ* studies ^1–5^. Despite extensive structural and functional characterization, the physiological significance of this supramolecular organization remains enigmatic. Respirasome formation has been proposed to optimize electron transfer, limit reactive oxygen species production, facilitate respiratory chain assembly and stabilize its constituent complexes ^6–8^. The bioenergetic significance of respirasome formation has received considerable attention but remains uncertain *in vivo* ^9^, raising the possibility that this supramolecular organization serves other physiological functions. One largely unexplored possibility is that respirasomes support mitochondrial proteostasis by facilitating respiratory chain assembly, stabilizing constituent complexes and/or limiting unfavorable protein interactions within the densely packed inner mitochondrial membrane.

Early evidence for a role of respirasomes in respiratory chain complex maintenance came from studies showing that defects in one complex can compromise another. Loss or severe impairment of CIII_2_ was shown to cause secondary depletion of CI, while defects in CIV have also been linked to impaired CI assembly or stability ^10–12^. Such interdependence led to the proposal that association with CIII_2_ and CIV within respirasomes contributes to CI stability. More recent studies have shown that CI can associate with other respiratory complexes before its assembly is complete, while CIV can undergo its final maturation within the respirasome ^13–15^. These findings suggest that individual complex biogenesis and respirasome formation are closely intertwined. Consequently, when disruption of one complex causes the loss of another, it is difficult to distinguish whether this secondary loss results from impaired biogenesis or from instability after assembly. A distinct role for respirasomes in post-assembly complex maintenance therefore remains unresolved.

Selectively perturbing the interface between otherwise intact respiratory chain complexes provides an opportunity to separate respirasome organization from individual complex biogenesis. Disruption of the CI–CIII_2_ contact through the *Uqcrc1^DEL:E258–D260^* allele strongly depletes respirasomes while preserving assembly and function of the constituent complexes ^9^. This separation allows us to directly test whether incorporation into respirasomes contributes to the maintenance of these complexes after they have assembled. Such a contribution could manifest through altered protein stability or turnover, or as a proteostatic vulnerability that emerges when mitochondrial homeostasis is challenged.

Here, we combine complementary proteomic approaches that capture distinct dimensions of protein homeostasis across cellular and *in vivo* models to investigate respiratory chain proteostasis following respirasome depletion, both at baseline and under mitochondrial stress. In mouse embryonic fibroblasts (MEFs) with severe respirasome depletion, thermal proteome profiling (TPP) ^1^^6^ and pulse-SILAC ^17^ reveal minimal effects on protein thermal stability and turnover. Solubility proteome profiling across diverse mitochondrial perturbations further shows that severe depletion of respirasomes does not increase proteostatic vulnerability to acute stress. *In vivo*, we combined whole-body *Uqcrc1^DEL:E258-D260^* knockin with TFAM deletion in heart and skeletal muscle to examine respiratory chain maintenance during progressive impairment of OXPHOS biogenesis. Temporal proteome profiling across disease progression showed that TFAM deficiency induced similar proteome remodeling regardless of respirasome depletion, with no acceleration of respiratory chain protein loss. Together, our findings show that post-assembly maintenance of respiratory chain complexes is largely independent of their organization into respirasomes, across cellular and tissue contexts and under diverse mitochondrial challenges.

## Results

### Mitochondrial protein thermal stability and turnover are preserved despite severe respirasome depletion

We first investigated whether the profound loss of respirasomes alters the biophysical properties of the mitochondrial proteome. To address this question, we performed thermal proteome profiling (TPP) in wild-type (WT) and homozygous *Uqcrc1^DEL:E258-D260^* knockin (*Uqcrc1* KI) immortalized mouse embryonic fibroblasts (MEFs) cultured in glucose or galactose (Figure 1A), the latter growth condition increases the cellular reliance on mitochondrial oxidative metabolism. TPP measures proteome-wide changes in protein thermal stability, providing a sensitive readout of alterations in protein biophysical state in intact cells ^18,19^.

**Figure 1.**
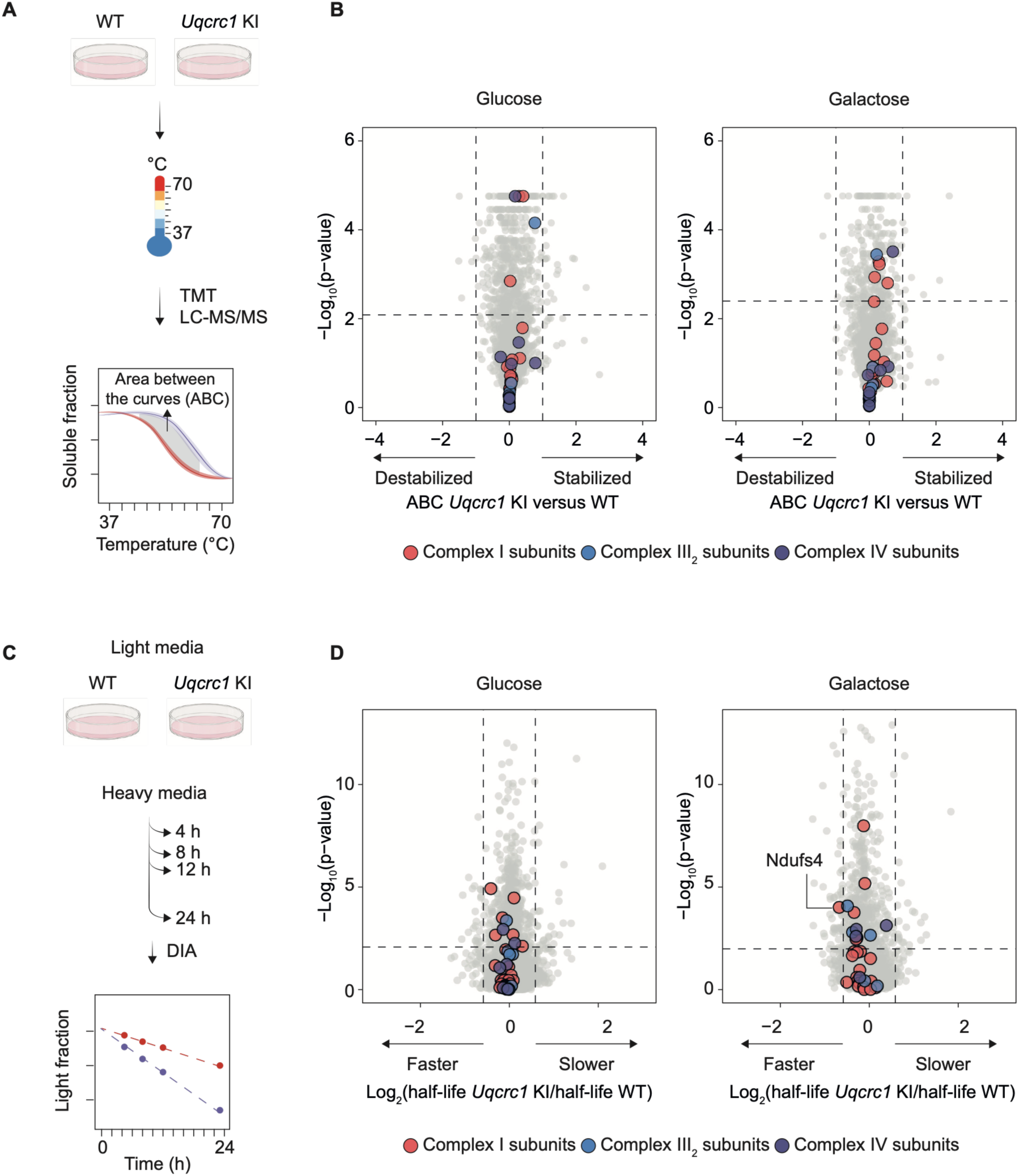
Thermal proteome profiling and pulse-SILAC reveal limited effects of respirasome depletion on protein stability and turnover in cells. (A) Schematic depiction of the thermal proteome profiling (TPP) workflow. Wild-type (WT) and homozygous *Uqcrc1^DEL:E258-D260^*knockin (*Uqcrc1* KI) immortalized mouse embryonic fibroblasts (MEFs) cultured in glucose or galactose were subjected to a temperature gradient (37–70°C), followed by TMT-based quantitative mass spectrometry and thermal stability analysis. (B) Volcano plots showing differences in thermal stability between WT and Uqcrc1 KI MEFs cultured in glucose or galactose. Subunits of respiratory chain complexes I, III_2_ and IV are highlighted as indicated. Positive and negative area between the curves (ABC) values indicate increased and decreased thermal stability in *Uqcrc1* KI relative to WT cells, respectively. The horizontal dashed line indicates the adjusted p-value threshold (< 0.05), and the vertical dashed lines indicate the ABC thresholds (±1). n=3 biological replicates per growth condition. (C) Schematic depiction of the pulse-SILAC workflow. WT and *Uqcrc1* KI MEFs cultured in glucose or galactose were switched from light to heavy SILAC medium and collected after 4, 8, 12, and 24 h. Heavy-label incorporation was quantified by DIA mass spectrometry, followed by protein turnover difference analysis. (D) Volcano plots showing differences in protein turnover between WT and *Uqcrc1* KI MEFs cultured in glucose or galactose. Subunits of complexes I, III_2_, and IV are highlighted as indicated. The horizontal dashed line indicates the adjusted p-value threshold (<0.05), and the vertical dashed lines indicate the log2 fold-change thresholds (±log2 (1.5)). n=3 biological replicates per growth condition.

Despite severe respirasome depletion in *Uqcrc1* KI MEFs, we detected no significant changes in thermal stability among the profiled subunits of the respirasome-forming complexes CI, CIII_2_ and CIV under either growth condition (Figure 1B). Likewise, we observed no coordinated changes in thermal stability across the mitochondrial proteome, irrespective of submitochondrial localization (Figure S1A).

Preserved thermal stability does not, however, exclude changes in mitochondrial protein turnover. Unassembled or destabilized proteins are typically recognized by quality-control pathways and undergo more rapid degradation ^20–23^. Thus, if respirasomes stabilize respiratory chain proteins, their loss may be reflected in altered protein turnover. To quantify protein turnover directly, we performed pulse stable isotope labeling by amino acids in cell culture (pulse-SILAC) in WT and *Uqcrc1* KI MEFs cultured in glucose or galactose. Cells were pulse-labelled for 4, 8, 12, and 24 h, and differences in protein turnover between WT and *Uqcrc1* KI cells were determined after correcting for cell proliferation (Figure 1C). Among the profiled CI, CIII_2_ and CIV subunits, NDUFS4 was the only protein that met the significance threshold, showing a modest ∼1.6-fold increase in turnover in *Uqcrc1* KI cells compared with WT cells, while turnover of the other profiled subunits remained unchanged (Figure 1D). At the broader mitochondrial level, we observed no differences in protein turnover across mitochondrial compartments (Figure S1B). Together, these findings provide no evidence for widespread destabilization or increased turnover of respiratory chain proteins following severe respirasome depletion.

### Profound loss of respirasomes does not compromise mitochondrial proteostasis under stress

The proteostatic consequences of severe respirasome depletion may become apparent only when mitochondrial homeostasis is challenged. If incorporation into respirasomes contributes to the stability of respiratory chain proteins, disruption of this organization could increase their susceptibility to stress-induced loss of solubility or aggregation. We therefore exposed WT and *Uqcrc1* KI MEFs to a panel of perturbations targeting distinct mitochondrial functions and assessed their proteostatic response by solubility proteome profiling ^24^ . Cells were treated for 16 h with compounds targeting iron homeostasis (deferiprone), membrane potential (CCCP), protein import (MitoBlock10), CIII_2_ (antimycin A), or ATP synthase (oligomycin) (Figure 2A). Following treatment, each sample was divided into two fractions. One fraction was extracted under mild detergent conditions and subjected to ultracentrifugation to quantify changes in the soluble proteome as a readout of protein aggregation, whereas the second was extracted under strong detergent conditions to quantify changes in total protein abundance. Across all stress conditions, we observed similar treatment-induced changes in mitochondrial proteins in WT and *Uqcrc1* KI cells in both the soluble and total proteomes (Figure 2B). Similarly, we found no genotype-dependent differences in the response of the profiled CI, CIII_2_ and CIV subunits to any of the stressors examined (Figure 2C). Thus, severe respirasome depletion does not increase susceptibility to stress-induced changes in protein solubility or abundance, neither in the respiratory chain nor across the broader mitochondrial proteome.

**Figure 2.**
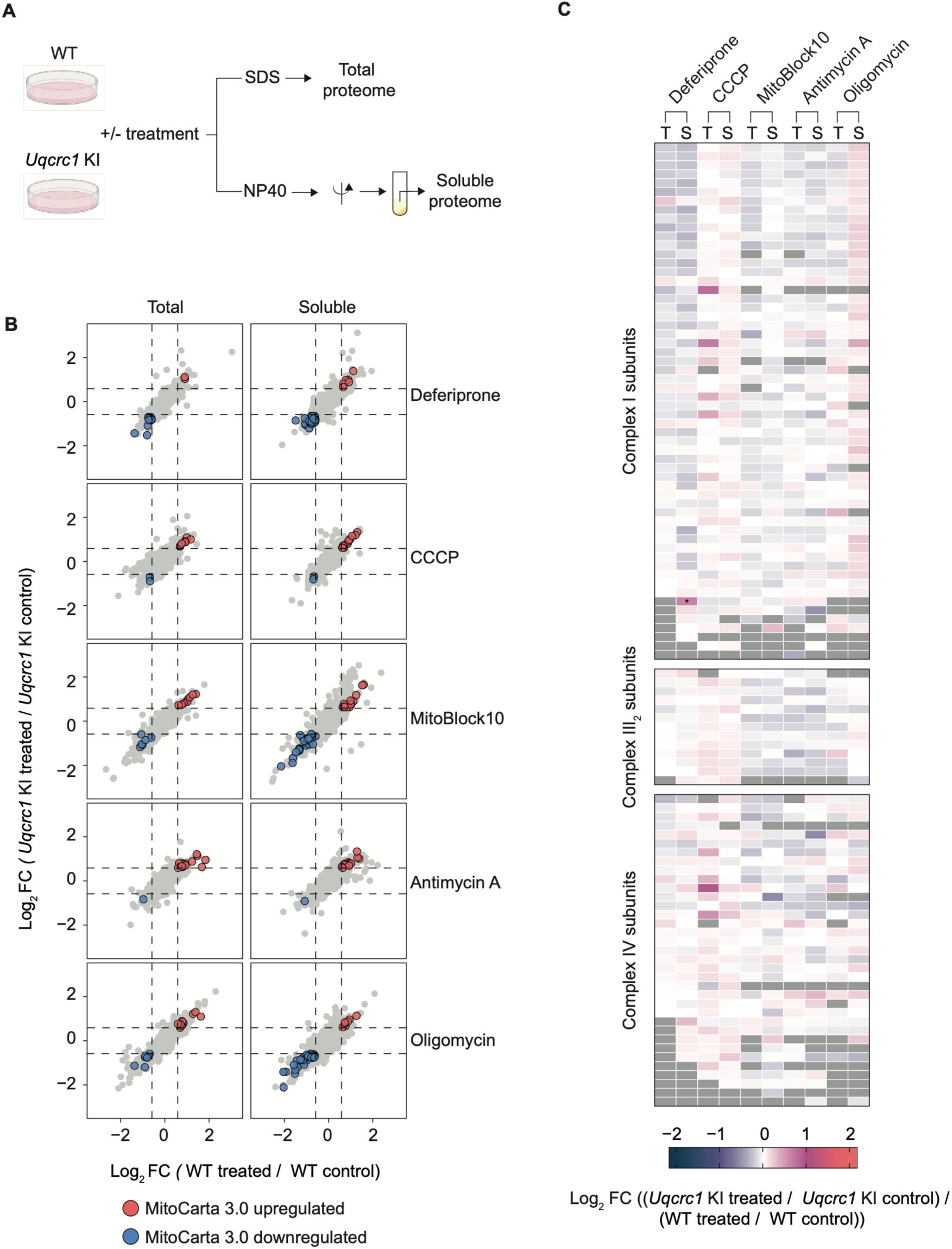
Respirasome depletion does not sensitize respiratory chain proteins to stress-induced solubility changes. **(A)** Schematic depiction of the solubility proteome profiling workflow. WT and *Uqcrc1* KI MEFs were exposed to the indicated mitochondrial stressors for 16 h. Samples were divided for extraction with either sodium dodecyl sulfate (SDS) to quantify the total proteome or NP-40 followed by ultracentrifugation to quantify the soluble proteome. **(B)** Scatter plots showing log2 fold changes in WT (WT treated versus WT control; x-axis) and *Uqcrc1* KI (KI treated versus KI control; y-axis) cells after treatment with each stressor in the total (left) and soluble (right) proteomes. Significantly upregulated and downregulated mitochondrial proteins (MitoCarta 3.0) are highlighted as indicated. Dashed lines indicate log2 fold-change thresholds (±log2 (1.5)). n=3 biological replicates per condition. (C) Heatmaps showing genotype-specific effects of the indicated mitochondrial stressors on complex I, III_2_ and IV subunits in the total (T) and soluble (S) proteomes. Gray denotes proteins that are not identified.

### TFAM deficiency induces similar cardiac remodeling with and without respirasome depletion

Given the preserved respiratory chain proteostasis observed in cells at baseline and under acute mitochondrial perturbations, we next asked whether requirement for respirasomes might emerge during mitochondrial dysfunction *in vivo*. To impose a progressive challenge to respiratory chain proteostasis, we crossed homozygous whole-body knockin *Uqcrc1^DEL:E258-D260^* mice ^9^ with conditional knockout mice carrying a heart- and skeletal muscle-specific deletion of a loxP-flanked allele for mitochondrial transcription factor A (*Tfam*) driven by cre-recombinase expressed from the muscle creatine kinase promoter (Ckmm-Cre) ^25,26^. TFAM is required for mitochondrial DNA (mtDNA) maintenance and transcription, and its loss ultimately impairs the biogenesis and function of the OXPHOS system ^25,27,28^. The resulting crosses yielded four experimental groups, *_TfamloxP/loxP Uqcrc1wt/wt_* _(WT), *TfamloxP/loxP Uqcrc1DEL:E258-D260/DEL:E258-D260 (Uqcrc1* KI), *TfamloxP/loxP;+/Ckmm-cre Uqcrc1wt/wt (Tfam* knockout; KO) and *TfamloxP/loxP;+/Ckmm-cre Uqcrc1DEL:E258-*_ *^D260/DEL:E258-D260^* (double mutant) (Figure 3A). *Tfam* KO and double-mutant mice showed closely overlapping survival, with both groups reaching the terminal disease stage at approximately 9 weeks of age (Figure 3B). Body weight was similarly affected in *Tfam* KO and double-mutant mice over the disease course (Figure 3C). Both groups developed a progressive increase in heart- to-body weight ratio, with no significant differences between *Tfam* KO and double-mutant mice at the examined time points, consistent with the development of cardiomyopathy (Figure 3D). To define disease progression of cardiac remodeling at the molecular level, we performed temporal proteomic profiling of hearts at 2, 3, 6 and 9 weeks of age (Figure 3E). Principal component analysis (PCA) revealed progressive remodeling of the cardiac proteome in *Tfam* KO hearts, with double-mutant hearts following a similar age-dependent trajectory (Figure 3F). At the protein level, *Tfam* KO hearts showed induction of cardiac stress markers, including NPPA and ANKRD1, and fibrosis-associated proteins, including POSTN and LGALS3, accompanied by a broader decline in proteins involved in contractile function and Ca²⁺ handling (Figure 3G). These changes were closely recapitulated in double-mutant hearts, showing that respirasome depletion does not substantially alter the molecular progression of cardiac remodeling caused by TFAM deficiency.

**Figure 3.**
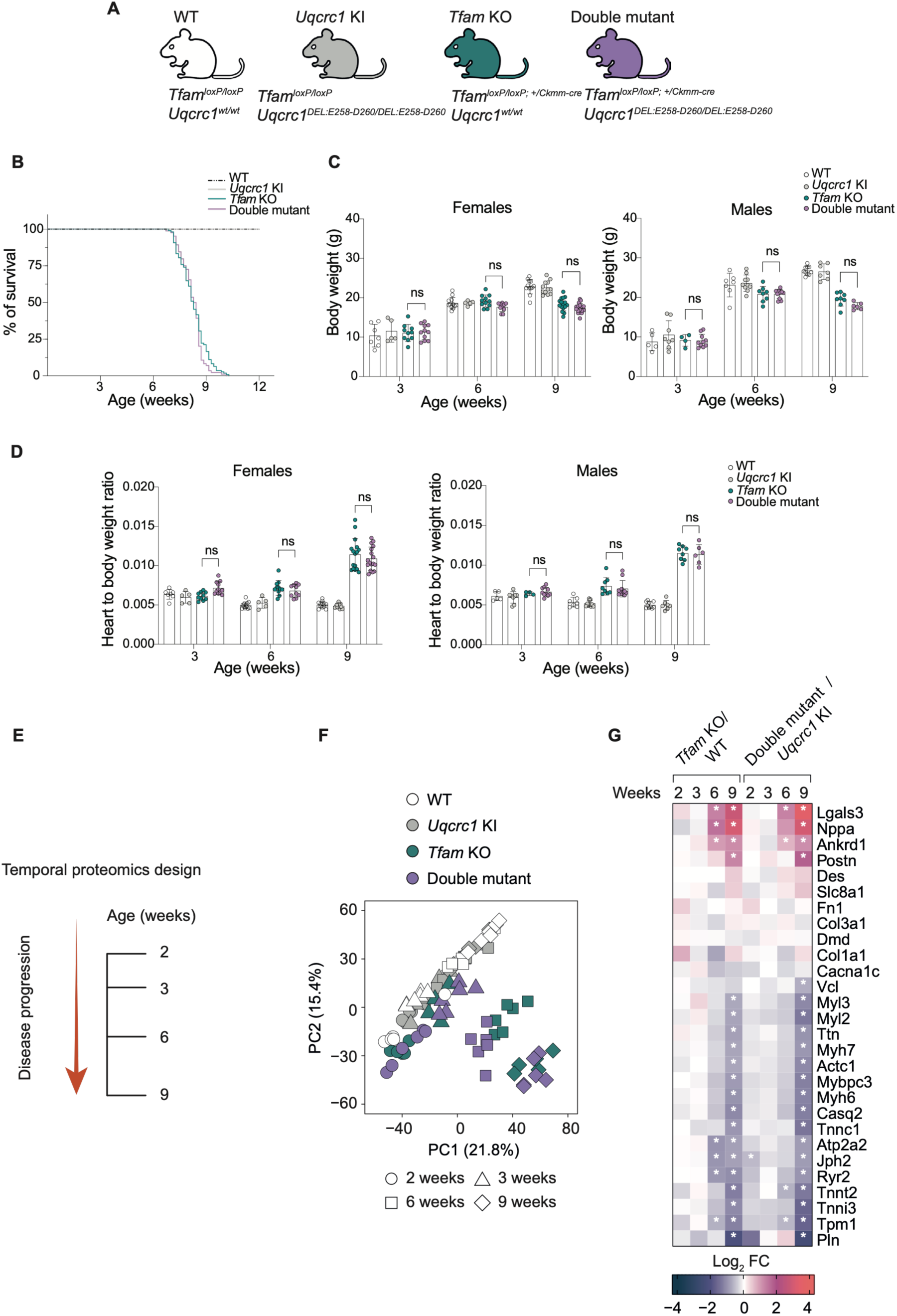
Temporal profiling reveals comparable cardiac remodeling following isolated TFAM deficiency or if combined with respirasome depletion. (A) Genotypes of the mouse models used to assess the effect of respirasome depletion during progressive mitochondrial dysfunction in the heart. Whole-body homozygous knockin *Uqcrc1^DEL:E258-D260^ (Uqcrc1* KI) mice were crossed with mice carrying a heart- and skeletal muscle-specific deletion of *Tfam* driven by *Ckmm-Cre*, generating WT, *Uqcrc1* KI, *Tfam* KO and double-mutant mice. (B) Kaplan–Meier survival curves of WT, *Uqcrc1* KI, *Tfam* KO and double-mutant mice. Median survival was 58 days for *Tfam* KO and 58.5 days for double-mutant mice. A total of 108 *Tfam* KO and 84 double-mutant mice were included in the survival analysis. (C) Body weight of female and male mice at 3, 6 and 9 weeks of age. (D) Heart-to-body weight ratio in female and male mice at 3, 6 and 9 weeks of age. Data are presented as mean ± SD with individual biological replicates. n=5-17 biological replicates per genotype and time point. Statistical significance was assessed using two-way ANOVA with genotype and age as factors, including the genotype × age interaction, followed by Tukey’s multiple-comparisons test comparing genotypes within each age. Brackets indicate comparisons between *Tfam* KO and double-mutant mice. ns, not significant. (E) Schematic depiction of the temporal proteomics design, with hearts collected at 2, 3, 6 and 9 weeks of age. **(F)** Principal component analysis (PCA) of the cardiac proteome from WT, *Uqcrc1* KI, *Tfam* KO and double-mutant mice at the indicated ages. Genotypes and ages are indicated by color and symbol, respectively. **(G)** Heatmap showing temporal changes in selected cardiac stress and remodeling proteins in *Tfam* KO and double-mutant hearts relative to their respective controls. Asterisks indicate significant changes (|log2 fold change| > log2 (1.5), adjusted p-value < 0.05). n=4-6 biological replicates per genotype and time point.

### Respirasome depletion does not accelerate respiratory chain protein loss during impaired OXPHOS biogenesis *in vivo*

A requirement for respirasomes in long-term complex maintenance could manifest as accelerated respiratory chain protein loss during progressive mitochondrial dysfunction. We therefore examined respiratory chain protein abundance over the disease course. TFAM deficiency induced similar changes in mitochondrial proteins in *Tfam* KO and double-mutant hearts at each time point, both in direction and magnitude (Figure 4A). Importantly, subunits of the respirasome-forming complexes CI, CIII_2_ and CIV underwent similar progressive changes in *Tfam* KO and double-mutant hearts, with no evidence that respirasome depletion accelerated their loss at any stage of disease progression (Figure 4B). A similar pattern was observed in skeletal muscle, profiled at 3, 6 and 9 weeks, where proteome changes emerged later and were less pronounced than in the heart. Nevertheless, respiratory chain proteins affected by TFAM deficiency showed comparable changes in *Tfam* KO and double-mutant skeletal muscle (Figure S2).

**Figure 4.**
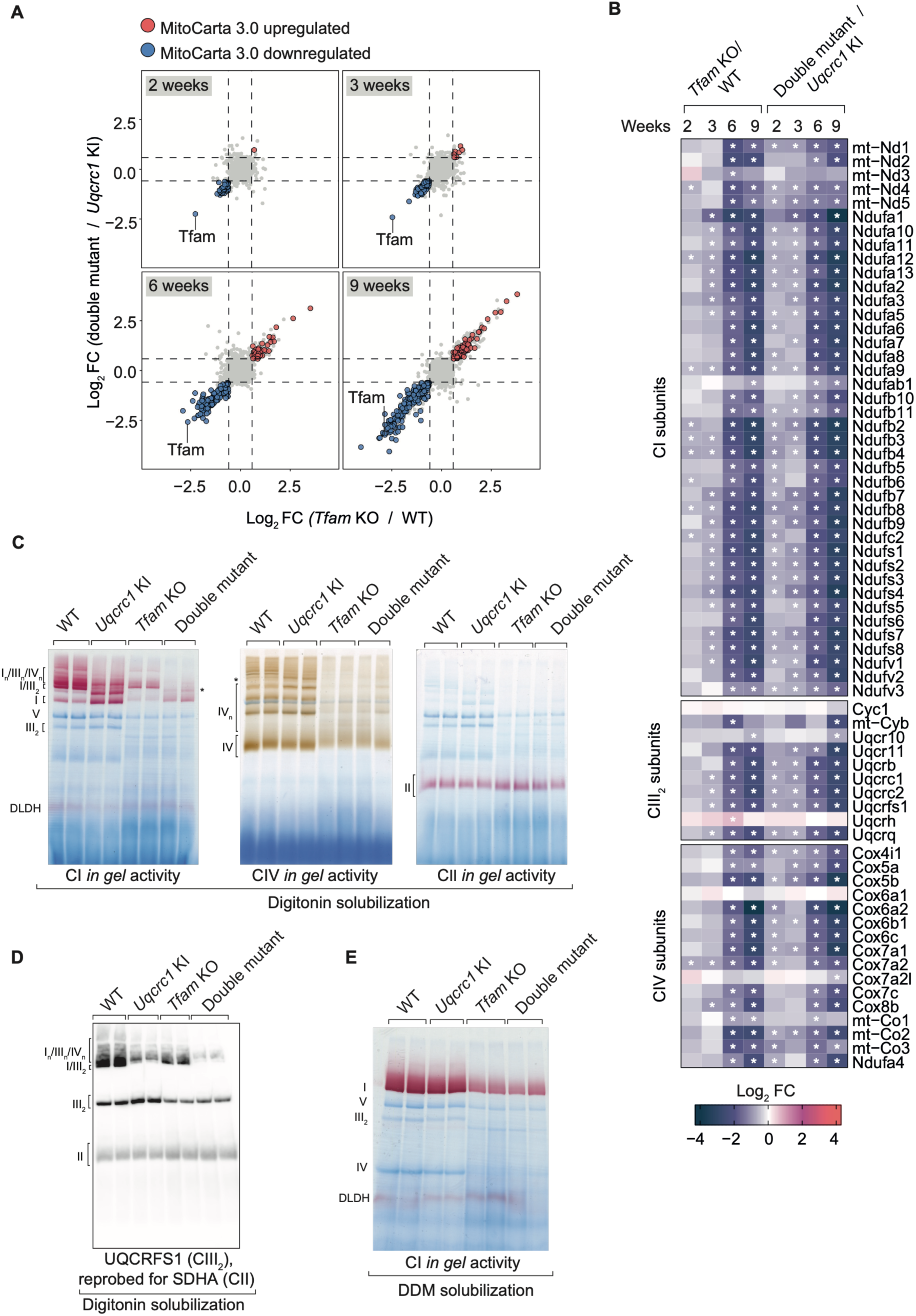
Progressive respiratory chain protein loss caused by TFAM deficiency is unaffected by respirasome depletion. **(A)** Scatter plots showing log2 fold changes in *Tfam* KO relative to WT (x-axis) and double-mutant relative to *Uqcrc1* KI (y-axis) at 2, 3, 6, and 9 weeks. Significantly upregulated and downregulated mitochondrial proteins (MitoCarta 3.0) are highlighted as indicated. Dashed lines indicate log2 fold-change thresholds (±log2 (1.5)). **(B)** Heatmaps showing log2 fold changes in protein abundance for CI, CIII_2_ and CIV subunits in *Tfam* KO versus WT and double-mutant versus *Uqcrc1* KI mice during disease progression. White stars indicate significant abundance changes. n=4-6 biological replicates per genotype and time point. **(C)** *In gel* activity assays of CI, CIV and CII following blue native PAGE (BN-PAGE) of digitonin-solubilized heart mitochondria from 6-week-old WT, *Uqcrc1* KI, *Tfam* KO and double-mutant mice. The asterisk indicates an aberrant supercomplex. **(D)** Western blot analysis of BN-PAGE-separated, digitonin- solubilized heart mitochondria using an antibody against UQCRFS1 (CIII_2_), reprobed with antibody against SDHA (CII) that serves as a loading control. **(E)** CI *in gel* activity following DDM solubilization of heart mitochondria from the indicated genotypes.

To assess respiratory chain organization, we analyzed heart mitochondria at 6 weeks of age by BN-PAGE. Following digitonin solubilization, *Uqcrc1* KI and double-mutant mitochondria showed severe respirasome depletion, accompanied by redistribution of CI and CIII_2_ into their free forms and accumulation of an aberrant supercomplex (Figure 4C-D and S3A) ^9^. Alongside the respirasome-disruption signature, double-mutant mitochondria showed a general decrease in respiratory chain complexes, similar to that observed in *Tfam* KO mitochondria and consistent with the downstream consequences of impaired mitochondrial gene expression. This general reduction in OXPHOS complexes was also evident by BN-PAGE analyses of n-dodecyl-β-D-maltoside (DDM)-solubilized heart mitochondria, which resolve the individual complexes and further confirmed that the *Uqcrc1^DEL:E25^*^8^*^-D260^* mutation alone does not impair their assembly (Figure 4E). Functional analyses of intact, respiring mitochondria provided further support for the absence of an additional respiratory chain defect in double-mutant hearts. Both CI- and CII-driven respiration were impaired by TFAM deficiency but were not further reduced in double-mutant mitochondria (Figure S3B). Similarly, H_2_O_2_ release from heart mitochondria was reduced in *Tfam* KO mitochondria, with no additional effect in double-mutant mitochondria (Figure S3C). Together, these findings show that severe respirasome depletion does not accelerate respiratory chain protein loss during progressive mitochondrial dysfunction, with no additional impairment in respiratory chain function.

## Discussion

Respiratory chain complex biogenesis and supercomplex formation were initially viewed as largely sequential processes, in which fully assembled individual complexes associate to form higher-order structures ^2^^9^. Increasing evidence, however, supports a more cooperative assembly model, in which individual complex biogenesis and supercomplex formation can proceed in parallel and influence one another ^13,30^. This interplay has made it difficult to distinguish a role for respirasomes during respiratory chain complex biogenesis from a requirement for their subsequent maintenance. By selectively disrupting respirasome formation while preserving individual complex assembly and function, our study distinguishes post-assembly maintenance from defects arising during complex biogenesis. Despite severe respirasome depletion, CI, CIII_2_ and CIV remained well maintained across cellular and tissue contexts, even under mitochondrial challenge. Thus, the post-assembly maintenance of individual complexes is largely independent of their organization into respirasomes.

Many respiratory chain proteins are long-lived, making mechanisms that preserve the integrity of assembled complexes over time particularly important ^31,32^. Studies in cells have suggested that supercomplex organization may contribute to this longevity, with supercomplex-associated subunits displaying slower turnover than those residing in individual complexes ^32^.

However, severe respirasome depletion did not substantially alter respiratory chain protein stability, turnover or solubility in cells. A requirement for respirasomes in complex maintenance could nevertheless become more apparent *in vivo*, where long-term metabolic labeling studies have shown that subsets of OXPHOS proteins can persist for months in tissues such as brain and heart ^33^. TFAM deficiency in heart and skeletal muscle provided a stringent context for examining complex maintenance, as impaired mitochondrial gene expression leads to a progressive decline in OXPHOS biogenesis over the course of disease. As the capacity to replace complexes becomes increasingly limited, any stabilizing contribution of respirasomes could become evident as accelerated protein loss. Yet severe respirasome depletion did not accelerate the loss of CI, CIII_2_ or CIV subunits in either tissue.

The maintenance of individual complexes is supported by mitochondrial quality-control mechanisms operating at multiple levels. During respiratory chain biogenesis, subunits that fail to assemble are rapidly targeted for proteolytic clearance rather than allowed to accumulate ^22,34^. This surveillance can begin as early as mitochondrial translation, where newly synthesized mtDNA-encoded proteins are subject to molecular triage at the mitoribosome and directed toward either productive assembly or proteolytic clearance ^35^. Quality control also extends beyond initial assembly. Within mature CI, damaged N-module components can be selectively removed and replaced without degradation of the entire complex ^36^. Similarly, defective COX1 can be incorporated into fully assembled CIV and even supercomplexes but remains subject to proteolytic clearance ^37,38^, suggesting that higher order assemblies do not necessarily shield respiratory chain proteins from quality-control surveillance. Together, these studies show that respiratory chain proteostasis is actively maintained at multiple stages of complex biogenesis and throughout the lifetime of assembled complexes. Our findings now show that respiratory chain complex maintenance is preserved even when higher-order organization into respirasomes is severely disrupted. This argues against a major requirement for respirasomes as stabilizing scaffolds.

The organization of individual complexes into respirasomes may reflect principles distinct from the functions ultimately conferred by these assemblies. Recent work supports this possibility by showing that formation of the CI–CIII_2_ supercomplex can alleviate protein-induced membrane strain and may therefore be favored by the physical properties of the inner mitochondrial membrane ^39^. Our findings further refine this distinction by showing that, once complexes are assembled, their maintenance is largely preserved despite disruption of respirasome organization, across both acute cellular perturbations and prolonged mitochondrial dysfunction in post-mitotic tissues. The functional significance of respirasomes may therefore lie in more context-dependent roles that emerge under specific cellular or physiological demands on the respiratory chain. Defining these contexts will be important for understanding when and how respirasome organization confers a physiological advantage.

## Acknowledgements

NGL was supported by the Swedish Research Council (2025-02712), the Swedish Cancer Foundation (243520Pj), the Knut and Alice Wallenberg Foundation (2023.0224 and 2024.0081), the Swedish Brain Foundation (FO2025-0061-HK-269), the Swedish Diabetes Foundation (DIA2025-952), the European Research Council (Advanced Grant 101141290), the Novo Nordisk Foundation (NNF25OC0105033), and by grants provided by Region of Stockholm (ALF project). MS was supported by the Allen Distinguished Investigator award through the Paul G. Allen Frontiers Group. PRM was supported by a Bridging Excellence Fellowship from the EMBL/Stanford Life Science Alliance.

## Author contributions

Conceptualization: JM, PRM, DM, MS and NGL; Data curation: JM and PRM; Formal analysis: JM and PRM; Funding acquisition: NGL and MS; Investigation: JM, PRM, CUN, NH and PM; Methodology: JM, PRM, IA and XL; Visualization: JM and PRM; Writing - original draft: JM, PRM, MS and NGL; and Writing - review & editing: all authors.

## Materials and Methods

### Mouse embryonic fibroblast isolation and immortalization

Primary mouse embryonic fibroblasts (MEFs) were isolated from individual embryos at embryonic day 13.5–14.5 (E13.5–E14.5) obtained from heterozygous *Uqcrc1^DEL:E258-D260^* intercrosses. Following removal of the head and internal red organs, embryonic tissue was finely minced and dissociated in TrypLE Express (Gibco) for 30 min at 37 °C. Digestion was stopped by addition of MEF medium, and the cell suspension was further mechanically dissociated and plated in DMEM supplemented with 10% fetal bovine serum (Gibco, 10270) and 1% penicillin–streptomycin. The head from each embryo was retained for genotyping. Primary MEFs were expanded and cryopreserved at passage 0.

For immortalization, early-passage MEFs (P1–P3) were transiently transfected with an SV40 plasmid using Lipofectamine 3000 (Thermo Fisher Scientific). Following transfection, cells were serially passaged at low density (1:10) to select for immortalized cells, and established MEF lines were subsequently expanded and cryopreserved.

## Cell culture

MEFs were maintained in Dulbecco’s Modified Eagle Medium (DMEM; Sigma-Aldrich, D5648) supplemented with 25 mM glucose or 10 mM galactose and 10% fetal bovine serum (Gibco, 10270). All cell lines were incubated at 37°C in a humidified atmosphere containing 5% CO2 and were routinely screened for Mycoplasma contamination.

For solubility proteome profiling, cells were maintained in Dulbecco’s modified Eagle medium (DMEM; Sigma-Aldrich, D5648) supplemented with 4.6 g/L glucose and 10% fetal bovine serum (Gibco, 10270) for 24 h, followed by treatment for 16 h with deferiprone (100 µM; Sigma-Aldrich, 379409), carbonyl cyanide m-chlorophenyl hydrazone (CCCP; 10 µM; Sigma-Aldrich, 215911), MitoBlock10 (10 µM; MedChemExpress, HY-115467), antimycin A (1 µM; Sigma-Aldrich, A8674), or oligomycin (1 µM; Sigma-Aldrich, O4876).

## Thermal proteome profiling and lysis

MEFs were washed and collected in PBS, and ten 20 μL aliquots of the cell suspension (1 × 10⁶ cells per temperature) were transferred into a 96-well PCR plate. The samples were centrifuged at 100 g for 30 seconds and heated for 3 minutes at temperatures ranging from 37°C to 67°C using a PCR machine (Agilent SureCycler 8800). Samples were then equilibrated to room temperature for 3 minutes, flash-frozen in liquid nitrogen, and stored at −70°C until further analysis.

Frozen samples were thawed at 25 °C for 1 min and lysed by adding 30 µL of lysis buffer containing 0.5% NP-40, cOmplete protease inhibitor (Roche, 11873580001), PhosSTOP (Roche, 4906845001), 1.5 mM MgCl₂, and 0.25 U benzonase. Samples were resuspended by pipetting 20 times with a multichannel pipette and incubated for 1 h at 4 °C with shaking at 300 rpm. The plate was then centrifuged for 5 min at 4,000 × g and 4 °C, and aggregates were removed from the supernatant using a 0.45 µm pore-size 96-well filter plate (Millipore, MSHVN4550). The flow- through was transferred to a new plate, mixed 1:1 with 2× sample buffer (180 mM Tris, pH 6.8; 4% SDS; 20% glycerol; 0.1% bromophenol blue), and stored at −20 °C.

## Solubility proteome profiling and lysis

MEFs treated with the different stressors were resuspended in lysis buffer containing cOmplete protease inhibitors, PhosSTOP, 0.5% NP-40, 1.5 mM MgCl₂ in PBS. Samples were then divided in half. One half was centrifuged at 100,000 g at 4°C for 20 minutes to obtain supernatants containing the soluble protein pool. Both the supernatant and the second half of the lysed samples were incubated with and Benzonase (0.25 U/mL, Merck, 712206-3) followed by incubation with 1% SDS. Protein concentration was determined using the Rapid Gold BCA assay (Thermo Fisher Scientific, A55860). Equal sample amounts were used for multiplexed MS measurements.

## Digestion and labelling

The protein samples from TPP and SPP experiments were digested following a modified SP3 protocol ^40^. In brief, equal volumes of samples across temperatures (equivalent to 10 μg of proteins at the lowest temperature for TPP) were mixed with water to reach 20 μL and added to a bead suspension composed of a 1:1 mix of Sera-Mag A (Cytiva 45152105050250) and Sera-Mag-B (Cytiva 65152105050250) and 22 μL ethanol. This mix was incubated at room temperature for 15 minutes with shaking at 500 rpm. Next, the mix was placed into a 0.45 μm 96-well filter plate, and the bead-bound proteins were washed three times with 70% ethanol. Afterward, 40 μL of digestion solution (1.25 mM TCEP, 5 mM chloroacetamide, 0.2 μg trypsin/Lys-C mix (Promega, V5073), in 100 mM HEPES, pH 8.5) was added for protein digestion and incubated overnight at room temperature. The next day, peptides were eluted by centrifugation and vacuum-dried. Next, peptides were resuspended in 10 μL of water and labelled for 1 h at room temperature with 4 μL (∼11 μg/μL in acetonitrile) of TMT10plex or TMT18plex (Thermo Fisher Scientific). The reaction was quenched with 4 μL of 5% hydroxylamine. Experiments belonging to the same biological replicate were combined. Pooled samples were desalted by solid-phase extraction using an OASIS HLB μElution 30 μm plate (Waters, 186001828BA), washed twice with 100 μL 0.05% formic acid, eluted with 100 μL 80% acetonitrile.

## High-pH offline fractionation

Dried peptide of TPP experiments samples were resuspended in 17 μL of 20 mM ammonium formate, pH 10, and fractionated into 48 fractions using C18-based reversed-phase chromatography with a Phenomenex Gemini 3-μm C18 110-Å 100-mm × 1-mm column. The samples were resolved over an 85-minute gradient (mobile phase A: 20 mM ammonium formate [pH 10]; mobile phase B: acetonitrile) at 0.1 mL/min, starting at 0% B, followed by a linear increase to 35% B from 2 to 60 minutes, a subsequent increase to 85% B from 62 to 68 minutes, followed by a linear decrease to 0% B up to 70 minutes, and finishing with equilibration of the system at 0% B until the end of the run. Finally, fractions were concatenated into 12 pools, and vacuum-dried.

## PGC offline fractionation

Dried peptides from SPP experiments were resuspended in 18 µL of buffer A (0.1% TFA and 2% ACN in MS-grade H₂O), and 15 µL was injected for fractionation. Peptides were separated by porous graphitic carbon (PGC) chromatography using a Hypercarb column (100 × 1.0 mm, 3 µm particle size; Thermo Fisher Scientific) maintained at 60 °C on an Ultimate 3000 LC system (Thermo Fisher Scientific) at a flow rate of 75 µL/min. Following a 1-min post-injection delay, the gradient started at 15% buffer B (0.1% TFA in ACN) and increased linearly to 45% B over 95 min, followed by an increase to 80% B over 5 min. The column was washed at 80% B for 5 min and subsequently re-equilibrated at 100% buffer A for 5 min. Fractions were collected from 4.5 to 100.5 min at 2-min intervals, yielding 48 fractions. The 48 fractions were concatenated into 12 pools and vacuum-dried.

## LC-MS DDA

Fractionated peptides were resuspended in 0.05% formic acid and separated using an UltiMate 3000 RSLC nano-LC system (Thermo Fisher Scientific) equipped with an analytical column (Waters nanoEase HSS C18 T3, 75 μm × 25 cm, 1.8 μm, 100 Å) and a trapping cartridge (precolumn; C18 PepMap 100, 5 μm, 300-μm i.d. × 5 mm, 100 Å). The LC system was directly coupled to a Q Exactive Plus mass spectrometer (Thermo Fisher Scientific) using a Nanospray-Flex ion source and a PicoTip Emitter (360-μm o.d. × 20-μm i.d.; 10-μm tip, New Objective). The mobile phase consisted of 0.1% formic acid in LC–MS-grade water (buffer A) and 0.1% formic acid in LC–MS-grade acetonitrile (buffer B). Peptides were loaded onto the trapping cartridge using a flow of 30 μL/min of solvent A for 3 minutes and eluted with a constant flow of 0.3 μL/min using 120 minutes of analysis time. During the elution step, the percentage of solvent B was increased stepwise: 2% to 4% B in 4 min, 4% to 8% in 2 min, 8% to 28% in 96 min, and 28% to 40% in another 10 min. A column-cleaning step using 80% B for 3 min was applied before the system was returned to its initial conditions (2% B) for re-equilibration for 10 minutes. The mass spectrometer was operated in positive ion mode with a spray voltage of 2.2 kV and a capillary temperature of 275°C. Full-scan mass spectrometry spectra with a mass range of 375–1,200 m/z were acquired in profile mode using a resolution of 70,000 (maximum fill time of 250 ms or a maximum of 3 × 10⁶ ions [automatic gain control, AGC]). The instrument cycled between MS and MS/MS acquisition in a data-dependent mode and consecutively fragmented the top 10 peaks with charges 2–4 on the MS scan. MS/MS spectra were acquired in profile mode in the Orbitrap with a resolution of 35,000, a maximum fill time of 120 ms, and an AGC target of 2 × 10⁵ ions, with a 30-s dynamic exclusion window (normalized collision energy was 30).

## Isotopic labelling

MEFs were cultured in SILAC DMEM Flex Media (Gibco, A2493901) containing “Light” (12C- and 14N-labelled, Arg0 and Lys0, Silantes, 211004102) arginine and lysine, 25 mM glucose or 10 mM galactose, 10% dialyzed FBS (Gibco, 26400044), and 1 mM L-glutamine. Cells were grown for 6 doublings before starting the time course. For the SILAC time course, cells were seeded onto 6-well plates and cultured for 24 h before medium exchange. Then, cells were incubated in media containing “Heavy” (13C- and 15N-labelled, Arg10 and Lys8, Silantes, 201604102) arginine and lysine, 25 mM glucose or 10 mM galactose, 10% dialyzed FBS, and 1 mM L-glutamine for 4, 8, 12, and 24 hours. Then, the medium was aspirated and cells were washed twice with warm PBS (with Ca and Mg), flash-frozen in liquid nitrogen, and stored at −70°C until further analysis.

## Lysis of SILAC-labelled samples

Frozen samples were thawed on ice, and 100 μL RIPA buffer (Thermo, 89901) supplemented with protease inhibitors (Roche, 11873580001) and 0.25 U benzonase (Merck, 71206-3) was added. Samples were incubated for 1 hour at 4°C with shaking at 300 rpm. Then, SDS was added to the samples to a final concentration of 2%, and samples were centrifuged at 160000 g for minutes to discard insoluble material. Then, samples were stored at −80°C until further analysis.

## Digestion of SILAC-labelled samples

The protein samples were digested following a modified SP3 protocol ^40^. In brief, 10 μg of proteins per sample were mixed with water to reach 20 μL and added to a bead suspension composed of a 1:1 mix of Sera-Mag A (Cytiva, 45152105050250) and Sera-Mag-B (Cytiva, 65152105050250), 30 μL ethanol, and 10 μL 15% formic acid. This mix was incubated at room temperature for 15 minutes with shaking at 500 rpm. Next, the mix was placed into a 0.45 μm 96-well filter plate, and the bead-bound proteins were washed three times with 70% ethanol. Afterward, 40 μL of digestion solution (1.25 mM TCEP, 5 mM chloroacetamide, 0.2 μg trypsin/Lys-C mix (Promega, cat, V5073), in 50 mM triethylammonium bicarbonate buffer, pH 8.5) was added for protein digestion and incubated overnight at room temperature. The next day, peptides were eluted by centrifugation, vacuum-dried, and stored at −20°C until MS analysis.

## LC-MS DIA of SILAC-labelled samples

Dried peptides were resuspended in 0.05% formic acid and separated using a Vanquish Neo UHPLC system (Thermo Fisher Scientific) operated in trap-and-elute mode. The LC was equipped with a trapping cartridge (precolumn; PepMap Neo C18, 5 μm, 300 μm inner diameter × 5 mm, 100 Å) and an analytical column (Ionopticks AUR3-25075C18-XT, 25 cm × 75 μm inner diameter, 1.7 μm C18). Solvent A comprised 0.1% formic acid supplemented with 3% DMSO in LC–MS-grade water, and solvent B comprised 0.1% formic acid supplemented with 3% DMSO in LC– MS-grade acetonitrile. Peptides were loaded onto the trapping cartridge and separated on the analytical column using a linear gradient of 5–26% B (flow rate of 300 nl min−1) for 29.7 min, followed by an increase to 40% B within 3 min before washing at 85% B for 5 min and re-equilibration to initial conditions (total MS acquisition time of 40 min). The LC system was coupled to an Orbitrap Astral mass spectrometer (Thermo Fisher Scientific) operated in DIA mode. The instrument was operated in positive ion mode with a spray voltage of 1.8 kV and a capillary temperature of 280°C. Full-scan MS spectra were acquired in the Orbitrap at a resolution of 120,000 with a mass range of 380-880 m/z, an AGC target of 5 × 10⁶ charges, and a maximum injection time of 5 ms. DIA spectra were acquired in the Astral mass analyzer with 4 m/z windows between 380-880 m/z. The MS2 scan range was set to 150–2,000 m/z, the normalized collision energy was set to 25%. The normalized AGC target was set to 500%, and the maximum injection time was set at 5 ms.

## Protein identification and quantification

For TPP and SPP experiments, mass spectrometry raw files were converted to mzML format using MSConvert (ProteoWizard) ^41^ with peak picking using the vendor’s algorithm. Files were searched using MSFragger101 v3.8 ^42^ in Fragpipe v20.0 against a database containing Mus musculus UniProt FASTA files (proteome ID UP000000589, downloaded on 09 March 2026), along with known contaminants and reverse protein sequences. The search parameters were as follows: tryptic digestion; maximum missed cleavages = 2; peptide length = 7–50; precursor mass tolerance = 20 ppm; fragment mass tolerance = 20; minimum peaks = 15; topN peaks = 300; fixed modifications were carbamidomethyl on cysteines and TMT18plex on lysine; variable modifications included acetylation on protein N-terminus, oxidation of methionine, and TMT18plex on peptide N-termini. Peptide Spectrum Match (PSM) validation was performed using Philosopher version 5.1.0, and the false discovery rate was set to 1% at the PSM, peptide, and protein levels. For TMT quantification, unique and razor peptides were used. Protein or peptide TSV files were processed using the R programming language.

For pulse-SILAC DIA analysis, raw files were analyzed using DIA-NN (v.2.1.0) ^43^ using an in silico DIA-NN-predicted spectral library. The spectral library was generated using Trypsin specificity, allowing one missed cleavage, with carbamidomethylation of cysteine as a fixed modification, oxidation of methionine as a variable modification and N-terminal methionine excision enabled. Peptide lengths of 6–30 amino acids, precursor charges of 1–4 and a precursor m/z range of 380–880 were considered. The spectral library was generated from fasta files downloaded from UniProt (Mouse: proteome ID UP000000589, downloaded on 22 January 2025). The DIA-NN search used following parameters: precursor FDR (%) = 1, legacy quantification, MBR enabled, and RT-dependent cross-run normalization. The channel-based search using the plexDIA framework ^44^ was then performed to quantify heavy-to-light SILAC ratios. It was configured using the following additional parameters: --fixed-mod SILAC,0.0,KR,label, --lib-fixed-mod SILAC, --channels SILAC,L,KR,0:0; SILAC,H,KR,8.014199:10.008269, --original-mods and --channel-run-norm. The report.parquet output was used for the downstream analysis.

## Data preprocessing and thermal stability analysis

Proteins detected in at least two replicates and with more than one unique peptide were retained for further analysis. Data preprocessing began by aligning raw intensities across MS runs, selecting the lowest temperature per condition in each experiment, and calculating the geometric mean for each protein. A scaling value was then determined as the ratio of the raw intensity at the lowest temperature per condition to the geometric mean of the raw intensities at all temperatures. All intensities were adjusted by multiplying the raw intensities by the scaling value for each MS run. Next, quantile normalization was applied to the aligned intensities, followed by scaling the normalized intensities by dividing by the replicate mean. This scaling method was preferred since it has been shown to be more robust to technical errors, improving the reproducibility of replicates within each condition.

The GP-Melt method was used to analyze melting curves and detect hits, as previously described ^45^. This approach models the data using a hierarchical structure: the top level represents the protein across all conditions and replicates, the second level corresponds to experimental conditions, and the third level accounts for replicates within each condition. Model fitting yields a predictive distribution for the melting curves. For visualization, melting curves are displayed as the mean and 95% confidence interval (CI) of the predictive distribution. To identify hits, a null model is defined by merging the two conditions into a single hypothetical condition. A likelihood ratio test (LRT) is performed by comparing the likelihoods of the alternative and null models. P-values are derived by approximating the null distribution of the LRT statistic, achieved by generating ten samples per protein from the null model, fitting the three-level hierarchical GP model to these samples, and aggregating the resulting LRT values. This process defines p-values similarly to permutation p-values. All p-values were adjusted using the Benjamini-Hochberg (BH) procedure. The area between curves (ABC) was calculated from the posterior mean of the predicted distributions after excluding the two lowest and two highest temperatures. Proteins with |ABC| > 1, corresponding to an estimated ∼2 °C shift in apparent melting temperature, and a Benjamini–Hochberg-adjusted p-value < 0.05 were considered significant.

## Data processing for solubility profiling

Solubility-profiling raw data were first filtered to retain proteins quantified with more than two unique peptides and were subsequently normalized using variance-stabilizing normalization (VSN). Log2-transformed normalized signal intensities from the NP-40 and SDS fractions were analyzed using the limma package to compare treated versus untreated samples within each genotype and to assess genotype-dependent treatment effects using the contrast (*Uqcrc1* KI treated/*Uqcrc1* KI control)/(WT treated/WT control). Proteins with |log2(FC)| > 0.5 and a Benjamini–Hochberg-adjusted P value < 0.05 were considered significant.

## Data processing for pulse-SILAC

### Protein turnover analysis

All analysis was done in R based on the report.parquet DIA-NN output file. Precursors were filtered for Lib.Q.Value < 0.01 and the Ms1.Normalised precursor intensities from matched light pL and heavy pH precursors were used to calculate the fraction of the light precursor f_L_:

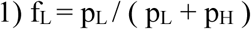

The light fraction of a protein FL was then calculated from the mean precursor Fraction L for each time point and replicate with at least 5 values and only considering values within the inner 80% quantile.

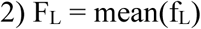

Each protein was then filtered per experimental group based on a min. of two data points per time point in at least two consecutive time points. The protein half-life t1/2 was then derived from an exponential decay model ^46^ fitted via the R lm() function for each protein in each experimental group separately:

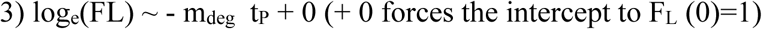

The apparent protein half-life T_1/2_,app is then given by

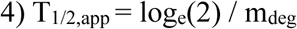

Cell cycle times were estimated based on the 100 proteins with the slowest turnover, for which the light-to-heavy ratio should only change due to cell growth. We selected the 100 proteins with the longest estimated half-live after filtering for R2 > 0.95 and min. 11 data points. Using the mean of the 100 longest half-lives Tcc, we corrected the FL to the fraction of old protein F_Old_ via

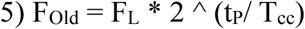

Values above 1 were set to 1 but subsequently not used to determine the protein half-lives. The corrected protein half-lives T_1/2_ were then calculated from F_Old_ instead of FL as described in (3) and (4).

### Comparative protein turnover fitting

To identify protein half-lives that change between genotypes withing each culture condition, we employed the F-statistic approach to derive a measure of how much the protein turnover profiles differed ^47^. Protein decay profiles of both groups were fitted together (H0 model) or separately (H1 model) as described above and the calibrated F-statistic distribution was derived by estimating the effective degrees of freedom for six separate bins depending on the number of total data points for each protein. The p-values were then calculated using the R pf() function. As a measure of effect size, we calculated the log2 fold change in protein half-life in *Uqcrc1* KI relative to WT cells [log2(half-life *Uqcrc1* KI/ half-life WT)]. Proteins with |log2(FC)| > 0.5 and a Benjamini– Hochberg-adjusted p-value < 0.05 were considered significant.

## Animals and housing

Heart and skeletal muscle-specific *Tfam* knockout mice (*Tfam^loxP/loxP;^*^+/*Ckmm-Cre*^) and the homozygous whole body *Uqcrc1^DEL:E258-D260^* knockin mice were generated as previously described ^9,25^. These founder lines were crossed to generate double-mutant mice (*Tfam^loxP/loxP;^*^+/*Ckmm-Cre*^ *Uqcrc1^DEL:E258-D260/DEL:E258-D260^*). Littermate mice with the following genotypes were used as controls: wild-type (*Tfam^loxP/loxP^ Uqcrc1^wt/wt^)*, *Uqcrc1* knockin (*Tfam^loxP/loxP^ Uqcrc1^DEL:E258-^ _D260/DEL:E258-D260)_*_, and *Tfam* knockout (*TfamloxP/loxP;*+/*Ckmm-Cre Uqcrc1wt/wt*). Animals were_ maintained on a C57BL/6N genetic background and housed in standard individually ventilated cages under a 12 h light/12 h dark cycle with *ad libitum* access to standard chow diet and water. Both male and female mice were included unless otherwise indicated. All animal experiments were approved by the appropriate animal welfare ethics committee and performed in accordance with national legislation and European Union guidelines.

## Sample preparation for label-free quantitative analysis of heart and skeletal muscle

Heart tissues were collected from mice aged 2, 3, 6, and 9 weeks, whereas quadriceps skeletal muscle tissues were collected from mice aged 3, 6, and 9 weeks. Approximately 10–20 mg of fresh tissue was cut into small pieces, snap-frozen in liquid nitrogen, and stored at −80 °C until further processing. Frozen tissues were homogenized using a TissueLyser III (QIAGEN) at a frequency of 20 Hz for 30–60 s until a homogeneous tissue powder was obtained. Subsequently, 50–100 µL of lysis buffer containing 6 M guanidinium chloride (GuHCl), 100 mM Tris-HCl, 2.5 mM tris(2- carboxyethyl)phosphine (TCEP), and 10 mM chloroacetamide (CAA) was added to each sample. The lysates were heated at 95 °C for 10 min, followed by sonication using a Bioruptor (Diagenode) for 10 min at high power (10 cycles of 30 s on and 30 s off). After centrifugation at 20,000 × g for 20 min, the supernatants were transferred to clean tubes. Protein concentrations were determined using an Epoch 2 microplate spectrophotometer (BioTek/Agilent). An aliquot containing 100 µg of protein from each sample was transferred to a 96-well deep-well plate, and the volume was adjusted to 10 µL using lysis buffer containing 6 M GuHCl. Proteins were subsequently processed using MagReSyn® Hydroxyl magnetic beads (ReSyn Biosciences), digested with trypsin (Promega), and the resulting peptides were purified using a KingFisher Apex automated purification system (Thermo Fisher Scientific).

## dia-PASEF analysis of heart and skeletal muscle

Peptides were separated using an Evosep One liquid chromatography system (Evosep) operated with the predefined 30 samples per day method. Chromatographic separation was performed on a ReproSil-Pur C18 analytical column (15 cm × 150 µm inner diameter, 1.9 µm particle size; EV1106 Endurance Column, Evosep). Mobile phases A and B consisted of 0.1% formic acid in water and 0.1% formic acid in acetonitrile, respectively. Peptides were analyzed using a trapped ion mobility spectrometry–quadrupole time-of-flight mass spectrometer (timsTOF Pro 2, Bruker) operated in data-independent acquisition parallel accumulation–serial fragmentation (diaPASEF) mode. The mass spectrometric acquisition range was set to m/z 100–1700, with TIMS ion accumulation and ramp times of 100 ms each. The ion mobility acquisition range was set to 1/K₀ = 0.85–1.30 V·s/cm². Each acquisition cycle consisted of one MS1 frame followed by 12 diaPASEF frames, resulting in a cycle time of approximately 1.38 s. The diaPASEF acquisition scheme comprised 24 isolation windows with variable isolation widths ranging from 15.5 to 43.1 Th, distributed across the m/z–ion mobility plane. Collision energy was varied linearly with ion mobility, using programmed collision-energy endpoints of 20–59 eV over a 1/K₀ range of 0.6–1.4 V·s/cm².

## Label-free proteomics data processing and quantification

Label-free raw LC-MS data were processed using Spectronaut version 18.5 (Biognosys) with default settings and the one-protein-per-gene reference proteome of Mus musculus (UniProt proteome ID: UP000000589; downloaded in August 2022). Methionine oxidation and protein N-terminal acetylation were specified as variable modifications, while cysteine carbamidomethylation was set as a fixed modification. Enzyme specificity was set to Trypsin/P, allowing up to two missed cleavages. Protein groups were retained if they contained at least n − 1 valid quantitative values in at least one experimental group, where n represents the total number of samples in that group. Missing values were imputed by random sampling from a down-shifted normal distribution (down-shift = 1.8; width parameter = 0.3). Differential protein abundance analysis was performed using the limma package version 3.34.9 ^48^ in R version 3.4.3 ^50^ .

## Mitochondria isolation

Heart mitochondria were obtained by differential centrifugation. Freshly excised hearts were washed in ice-cold phosphate-buffered saline (PBS), minced into small pieces and homogenized in mitochondrial isolation buffer (MIB; 310 mM sucrose, 10 mM Tris-HCl, pH 7.4, 1 mM EDTA, supplemented with freshly added 1× Complete protease inhibitor cocktail; Roche). The homogenate was first centrifuged at 1,000 × g for 10 min at 4°C to remove nuclei and tissue debris. Mitochondria were subsequently recovered by centrifugation of the supernatant at 10,000 × g for 15 min at 4°C. The resulting mitochondrial pellet was resuspended in an appropriate volume of MIB and used directly for subsequent experiments.

For high-resolution respirometry analyses and H_2_O_2_ measurements, heart mitochondria were isolated using a modified differential centrifugation protocol. Excised hearts were minced in ice-cold MIB (225 mM sucrose, 20 mM Tris-HCl, pH 7.4, 1 mM EDTA) and incubated with 2.5% trypsin (200 μl in 1.8 ml isolation buffer) for 10 min at 4°C. Trypsin activity was quenched by addition of MIB containing 0.04% trypsin inhibitor before the tissue was homogenized. Volume of the homogenate was adjusted to 10 ml and was centrifuged at 1,000 × g for 10 min at 4°C. The resulting supernatant was centrifuged at 10,000 × g for 10 min at 4°C. Obtained mitochondrial pellet was resuspended and washed sequentially with MIB containing 0.2% bovine serum albumin (BSA) and BSA-free MIB, with centrifugation (10,000 × g for 10 min at 4°C) after each wash. The final mitochondrial pellet was finally resuspended in BSA-free MIB and protein concentration was measured using a Qubit fluorometer (Invitrogen).

## BN-PAGE

Protein complexes were resolved by BN-PAGE as previously described ^49^. Mitochondrial proteins (75 µg) were extracted under native conditions by incubation on ice in 50 µl of solubilization buffer (20 mM Tris-HCl, pH 7.4, 0.1 mM EDTA, 50 mM NaCl, 10% glycerol) containing either 1–3% (w/v) digitonin (Calbiochem) or n-dodecyl-β-D-maltoside (DDM; Merck Millipore), as indicated. Solubilized samples were combined with Coomassie-based native sample buffer (5% Coomassie Brilliant Blue G-250, 150 mM Bis-Tris, 500 mM ε-amino-n-caproic acid, pH 7.0) and resolved on homemade 3–13% gradient BN-PAGE gels.

## Western blotting

Proteins were separated by BN-PAGE, transferred onto polyvinylidene difluoride (PVDF) membranes and analyzed by immunoblotting. Immunoreactive bands were visualized by enhanced chemiluminescence using Clarity Western ECL Substrate (Bio-Rad). Primary antibodies against UQCRFS1 (Abcam, ab14746) and SDHA (Abcam, ab14715) were used in this study.

### *In gel* activity assays

Complex I, II and IV activities were performed following the BN-PAGE by incubating gels in enzyme-specific reaction buffers. Complex I activity was detected in 2 mM Tris-HCl (pH 7.4) containing 0.1 mg/ml NADH (Roche) and 2.5 mg/ml iodonitrotetrazolium chloride (Thermo Scientific) for 5–10 min. For complex II activity, gels were incubated for 10–30 min in 5 mM Tris-HCl (pH 7.4) supplemented with 200 μl of 1 M sodium succinate, 8 μl of 250 mM phenazine methosulfate (prepared in DMSO), and 25 mg iodonitrotetrazolium chloride. Complex IV activity was determined by incubating gels for approximately 1 h in 10 ml of 0.05 M phosphate buffer (pH 7.4) containing 25 mg 3,3′-diaminobenzidine tetrahydrochloride (DAB), 50 mg cytochrome c, 3.75 g sucrose, and 1 mg catalase. Complex I and II activities were visualized by the formation of a red formazan precipitate, whereas complex IV activity was detected by DAB oxidation. All reactions were carried out at room temperature and terminated by incubation in a solution containing 50% methanol and 10% acetic acid.

## High-resolution respirometry analyses

Oxygen consumption by isolated heart mitochondria was measured using an Oxygraph-2k high-resolution respirometer (Oroboros Instruments, Innsbruck, Austria). A total of 30 µg of intact heart mitochondria was added to 2.1 mL of respiration buffer containing 120 mM sucrose, 50 mM KCl, 20 mM Tris-HCl, 4 mM KH₂PO₄, 2 mM MgCl₂, 1 mM EGTA, and 0.25 mg/mL fatty acid-free BSA (pH 7.2) at 37°C. Complex I-driven respiration was measured in the presence of 10 mM sodium pyruvate, 5 mM L-glutamate, and 5 mM L-malate. Complex II-driven respiration was measured using 10 mM succinate in the presence of 100 µM rotenone. For both substrate conditions, oxygen consumption was assessed in the phosphorylating state following the addition of 5 mM ADP, in the non-phosphorylating state following the addition of 0.5 µg/mL oligomycin, and at maximal uncoupled respiratory capacity following stepwise additions of 0.5 µM carbonyl cyanide m-chlorophenyl hydrazone (CCCP). Oxygen consumption was continuously recorded throughout the experiment using DatLab software (Oroboros Instruments, Innsbruck, Austria).

## H2O2 release by intact heart mitochondria

Hydrogen peroxide production was quantified in freshly isolated heart mitochondria using the Amplex Red Hydrogen Peroxide Assay Kit (Thermo Fisher Scientific, A22188). Intact mitochondria (30 µg protein) were incubated at 37°C in respiration buffer supplemented with Amplex Red, horseradish peroxidase, succinate and oligomycin. Fluorescence was monitored continuously (excitation, 570 nm; emission, 590 nm). Rates of H_2_O_2_ production during succinate- supported respiration were determined from the linear increase in fluorescence and quantified using an H_2_O_2_ standard curve. Rotenone was subsequently added to inhibit complex I, allowing assessment of H_2_O_2_ generation associated with complex III-dependent electron transfer. H_2_O_2_ production rates were normalized to mitochondrial protein content and all samples were analyzed in duplicates.

## Statistical analyses

Sample sizes and statistical tests are indicated in the corresponding figure legends. All proteomics datasets were analyzed using R (version 4.2.2 and 3.4.3). Other statistical analyses and graph generation were performed using GraphPad Prism version 11. A p-value < 0.05 was considered statistically significant.

## Declaration of generative AI and AI-assisted technologies in the manuscript preparation process

During the preparation of this work, the authors used ChatGPT and Grammarly to assist with the writing of the manuscript. After using these tools, the authors reviewed and edited the content as needed and take full responsibility for the content of the published article.

## Data availability

The mass spectrometry-based proteomics data will be deposited with the ProteomeXchange Consortium via the PRIDE partner repository.

## Supplementary figures

**Figure S1.**
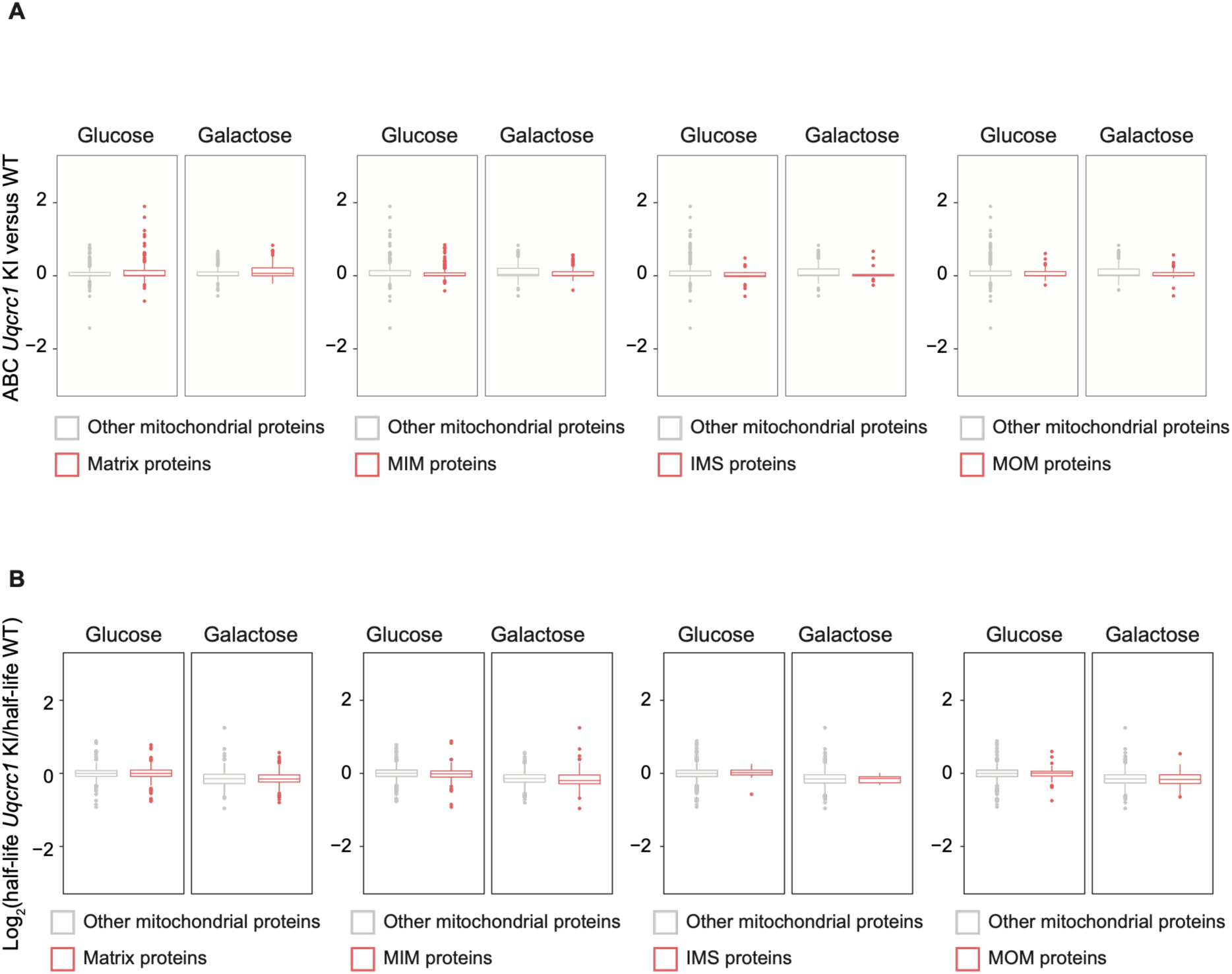
Respirasome depletion does not alter mitochondrial protein thermal stability or turnover across mitochondrial compartments. (A) Box plots showing thermal stability differences between *Uqcrc1* KI and WT MEFs cultured in glucose or galactose, grouped by submitochondrial localization. Proteins assigned to the mitochondrial matrix, inner mitochondrial membrane (MIM), intermembrane space (IMS), or mitochondrial outer membrane (MOM) are shown in red and compared with all other profiled mitochondrial proteins in gray. (B) Box plots showing the log2 ratio of protein half-life in *Uqcrc1* KI relative to WT MEFs cultured in glucose or galactose, grouped by the same submitochondrial compartments as in (A). Proteins assigned to each indicated compartment are shown in red and all other profiled mitochondrial proteins in gray. Boxes indicate the interquartile range, center lines indicate the median, whiskers extend to 1.5× the interquartile range, and points indicate outliers.

**Figure S2.**
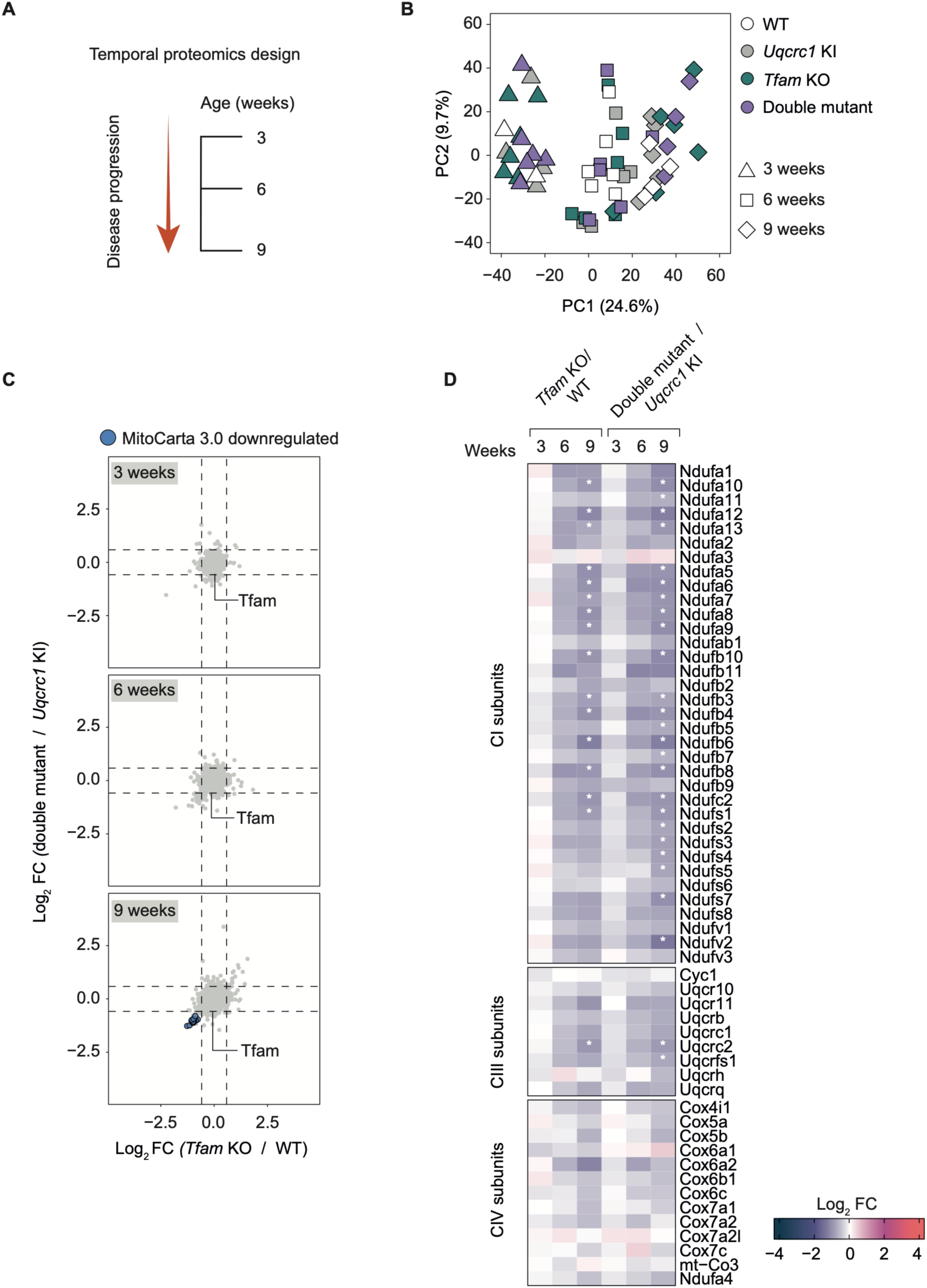
Skeletal muscle proteome responses to TFAM deficiency are largely unaffected by respirasome depletion. **(A)** Schematic depiction of the temporal proteomics design of skeletal muscle collected at 3, 6 and 9 weeks of age. **(B)** Principal component analysis (PCA) of skeletal muscle proteomes from WT, *Uqcrc1* KI, *Tfam* KO and double-mutant mice at 3, 6 and 9 weeks of age. Genotypes and ages are indicated by color and symbol, respectively. **(C)** Scatter plots comparing proteome changes in *Tfam* KO relative to WT (x-axis) and double mutant relative to *Uqcrc1* KI (y-axis) skeletal muscle at 3, 6 and 9 weeks of age. Significantly downregulated mitochondrial proteins (MitoCarta 3.0) are highlighted as indicated. Dashed lines indicate log2 fold-change thresholds (±log2(1.5)). **(D)** Heatmaps showing log2 fold changes in protein abundance for CI, CIII_2_, and CIV subunits in *Tfam* KO versus WT and double-mutant versus *Uqcrc1* KI mice during disease progression. White stars indicate significant abundance changes. n=5-6 biological replicates per genotype and time point.

**Figure S3.**
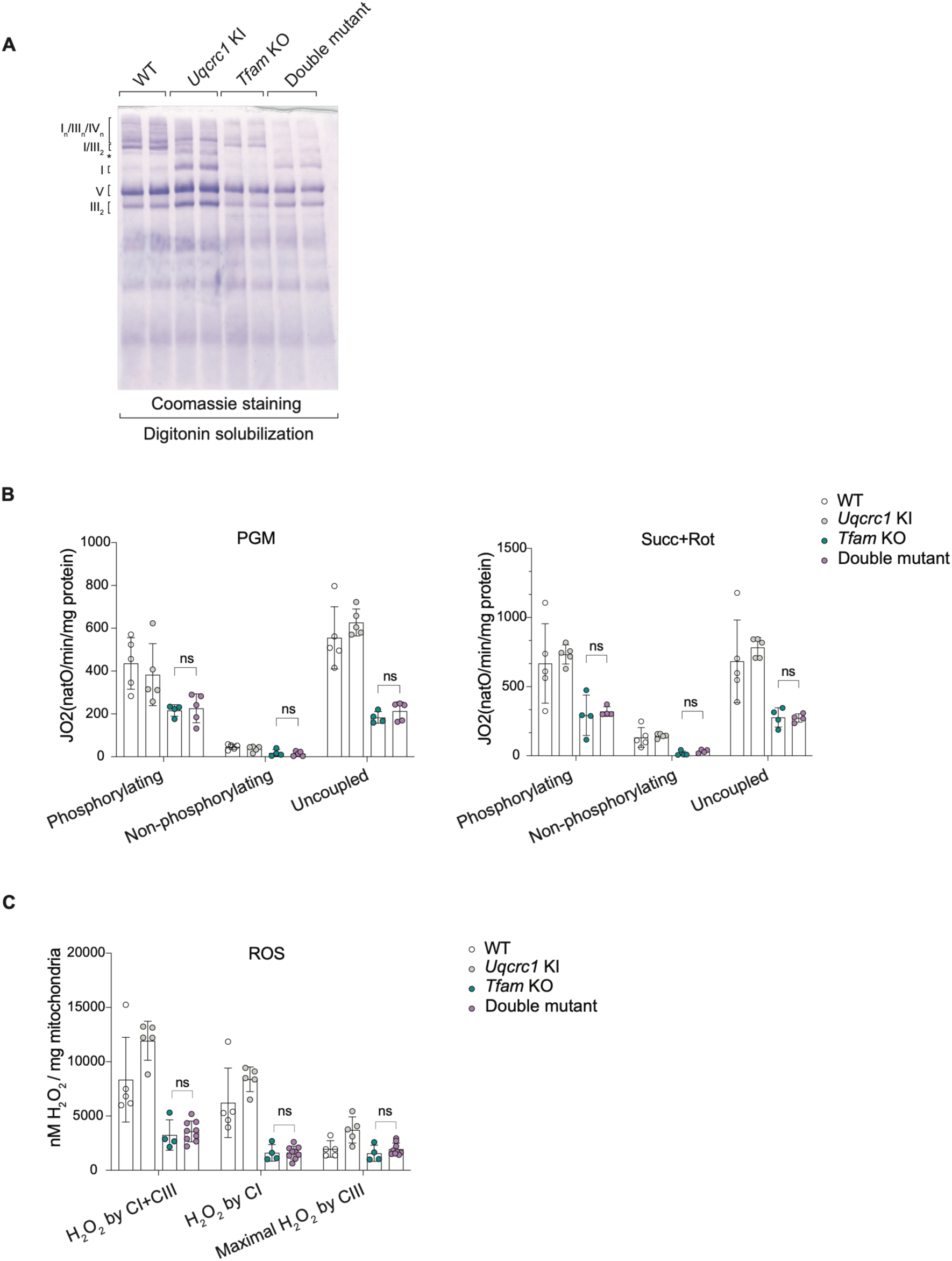
Respirasome depletion does not exacerbate respiratory chain dysfunction caused by TFAM deficiency in the heart. **(A)** Coomassie staining of BN-PAGE-separated, digitonin- solubilized heart mitochondria from 5-6-week-old WT, *Uqcrc1* KI, *Tfam* KO, and double-mutant mice. The asterisk indicates an aberrant supercomplex. **(B)** Oxygen consumption of intact heart mitochondria from 5–6-week-old mice of the indicated genotypes using pyruvate, glutamate and malate (PGM; left) or succinate and rotenone (Succ+Rot; right) as substrates. Phosphorylating, non-phosphorylating and uncoupled respiration measurements are shown. **(C)** H_2_O_2_ release from isolated heart mitochondria from 5–6-week-old mice. In both (**A)** and **(B)**, data are presented as mean ± SD with individual biological replicates. n=4-9 biological replicates per genotype. Statistical significance was assessed by two-way repeated-measures ANOVA with genotype and condition as factors, followed by Tukey’s multiple-comparisons test comparing genotypes within each condition. Brackets indicate comparisons between *Tfam* KO and double-mutant mice. ns, not significant.

## References

1. Gu, J., Wu, M., Guo, R., Yan, K., Lei, J., Gao, N., and Yang, M. (2016). The architecture of the mammalian respirasome. Nature 537, 639–643. 10.1038/nature19359.

2. Wu, M., Gu, J., Guo, R., Huang, Y., and Yang, M. (2016). Structure of Mammalian Respiratory Supercomplex I1III2IV1. Cell 167, 1598–1609.e10. 10.1016/J.CELL.2016.11.012.

3. Letts, J.A., Fiedorczuk, K., and Sazanov, L.A. (2016). The architecture of respiratory supercomplexes. Nature 537, 644–648. 10.1038/nature19774.

4. Vercellino, I., and Sazanov, L.A. (2024). SCAF1 drives the compositional diversity of mammalian respirasomes. Nat. Struct. Mol. Biol. 10.1038/s41594-024-01255-0.

5. Zheng, W., Chai, P., Zhu, J., and Zhang, K. (2024). High-resolution in situ structures of mammalian respiratory supercomplexes. Nature 2024, 1–8. 10.1038/s41586-024-07488-9.

6. Milenkovic, D., Blaza, J.N., Larsson, N.G., and Hirst, J. (2017). The Enigma of the Respiratory Chain Supercomplex. Preprint at Cell Press, 10.1016/j.cmet.2017.03.009.

7. Signes, A., and Fernandez-Vizarra, E. (2018). Assembly of mammalian oxidative phosphorylation complexes I-V and supercomplexes. Essays Biochem. 62, 255–270. 10.1042/EBC20170098.

8. Kohler, A., Barrientos, A., Fontanesi, F., and Ott, M. (2023). The functional significance of mitochondrial respiratory chain supercomplexes. EMBO Rep. 10.15252/EMBR.202357092.

9. Milenkovic, D., Misic, J., Hevler, J.F., Molinié, T., Chung, I., Atanassov, I., Li, X., Filograna, R., Mesaros, A., Mourier, A., et al. (2023). Preserved respiratory chain capacity and physiology in mice with profoundly reduced levels of mitochondrial respirasomes. Cell Metab. 35, 1799–1813.e7. 10.1016/j.cmet.2023.07.015.

10. Schägger, H., de Coo, R., Bauer, M.F., Hofmann, S., Godinot, C., and Brandt, U. (2004). Significance of respirasomes for the assembly/stability of human respiratory chain complex I. J. Biol. Chem. 279, 36349–36353. 10.1074/jbc.M404033200.

11. Acín-Pérez, R., Bayona-Bafaluy, M.P., Fernández-Silva, P., Moreno-Loshuertos, R., Pérez-Martos, A., Bruno, C., Moraes, C.T., and Enríquez, J.A. (2004). Respiratory complex III is required to maintain complex I in mammalian mitochondria. Mol. Cell 13, 805–815. 10.1016/s1097-2765(04)00124-8.

12. Li, Y., D’Aurelio, M., Deng, J.-H., Park, J.-S., Manfredi, G., Hu, P., Lu, J., and Bai, Y. (2007). An assembled complex IV maintains the stability and activity of complex I in mammalian mitochondria. J. Biol. Chem. 282, 17557–17562. 10.1074/jbc.M701056200.

13. Moreno-Lastres, D., Fontanesi, F., García-Consuegra, I., Martín, M.A., Arenas, J., Barrientos, A., and Ugalde, C. (2012). Mitochondrial complex I plays an essential role in human respirasome assembly. Cell Metab. 15, 324–335. 10.1016/j.cmet.2012.01.015.

14. Fang, H., Ye, X., Xie, J., Li, Y., Li, H., Bao, X., Yang, Y., Lin, Z., Jia, M., Han, Q., et al. (2021). A membrane arm of mitochondrial complex I sufficient to promote respirasome formation. Cell Rep. 35. 10.1016/j.celrep.2021.108963.

15. Nguyen, M.D., Sierra-Magro, A., Singh, V., Khawaja, A., Timón-Gómez, A., Barrientos, A., and Rorbach, J. (2026). Structural basis for late maturation steps of mitochondrial respiratory chain complex IV within the human respirasome. Nat. Commun. 17, 1550. 10.1038/s41467-025-68274-3.

16. Savitski, M.M., Reinhard, F.B.M., Franken, H., Werner, T., Savitski, M.F., Eberhard, D., Martinez Molina, D., Jafari, R., Dovega, R.B., Klaeger, S., et al. (2014). Tracking cancer drugs in living cells by thermal profiling of the proteome. Science 346, 1255784. 10.1126/science.1255784.

17. Schwanhäusser, B., Gossen, M., Dittmar, G., and Selbach, M. (2009). Global analysis of cellular protein translation by pulsed SILAC. Proteomics 9, 205–209. 10.1002/pmic.200800275.

18. Becher, I., Andrés-Pons, A., Romanov, N., Stein, F., Schramm, M., Baudin, F., Helm, D., Kurzawa, N., Mateus, A., Mackmull, M.-T., et al. (2018). Pervasive Protein Thermal Stability Variation during the Cell Cycle. Cell 173, 1495–1507.e18. 10.1016/j.cell.2018.03.053.

19. Mateus, A., Hevler, J., Bobonis, J., Kurzawa, N., Shah, M., Mitosch, K., Goemans, C. V, Helm, D., Stein, F., Typas, A., et al. (2020). The functional proteome landscape of Escherichia coli. Nature 588, 473–478. 10.1038/s41586-020-3002-5.

20. McShane, E., Sin, C., Zauber, H., Wells, J.N., Donnelly, N., Wang, X., Hou, J., Chen, W., Storchova, Z., Marsh, J.A., et al. (2016). Kinetic Analysis of Protein Stability Reveals Age-Dependent Degradation. Cell 167, 803–815.e21. 10.1016/j.cell.2016.09.015.

21. Song, J., Herrmann, J.M., and Becker, T. (2021). Quality control of the mitochondrial proteome. Nat. Rev. Mol. Cell Biol. 22, 54–70. 10.1038/s41580-020-00300-2.

22. Bogenhagen, D.F., and Haley, J.D. (2020). Pulse-chase SILAC-based analyses reveal selective oversynthesis and rapid turnover of mitochondrial protein components of respiratory complexes. J. Biol. Chem. 295, 2544–2554. 10.1074/jbc.RA119.011791.

23. Baker, M.J., Tatsuta, T., and Langer, T. (2011). Quality control of mitochondrial proteostasis. Cold Spring Harb. Perspect. Biol. 3, 1–19. 10.1101/cshperspect.a007559.

24. Sridharan, S., Kurzawa, N., Werner, T., Günthner, I., Helm, D., Huber, W., Bantscheff, M., and Savitski, M.M. (2019). Proteome-wide solubility and thermal stability profiling reveals distinct regulatory roles for ATP. Nat. Commun. 10, 1155. 10.1038/s41467-019-09107-y.

25. Larsson, N.G., Wang, J., Wilhelmsson, H., Oldfors, A., Rustin, P., Lewandoski, M., Barsh, G.S., and Clayton, D.A. (1998). Mitochondrial transcription factor A is necessary for mtDNA maintenance and embryogenesis in mice. Nat. Genet. 18, 231–236. 10.1038/ng0398-231.

26. Kühl, I., Miranda, M., Atanassov, I., Kuznetsova, I., Hinze, Y., Mourier, A., Filipovska, A., and Larsson, N.-G. (2017). Transcriptomic and proteomic landscape of mitochondrial dysfunction reveals secondary coenzyme Q deficiency in mammals. Elife 6, e30952. 10.7554/eLife.30952.

27. Kukat, C., and Larsson, N.-G. (2013). mtDNA makes a U-turn for the mitochondrial nucleoid. Trends Cell Biol. 23, 457–463. 10.1016/j.tcb.2013.04.009.

28. Rubalcava-Gracia, D., and Larsson, N.-G. (2026). Mammalian mtDNA gene expression: Concepts learned from in vivo models. Biochim. Biophys. Acta Mol. Cell Res. 1873, 120158. 10.1016/j.bbamcr.2026.120158.

29. Acín-Pérez, R., Fernández-Silva, P., Peleato, M.L., Pérez-Martos, A., and Enriquez, J.A. (2008). Respiratory active mitochondrial supercomplexes. Mol. Cell 32, 529–539. 10.1016/j.molcel.2008.10.021.

30. Fernández-Vizarra, E., and Ugalde, C. (2022). Cooperative assembly of the mitochondrial respiratory chain. Trends Biochem. Sci. 47, 999–1008. 10.1016/J.TIBS.2022.07.005.

31. Karunadharma, P.P., Basisty, N., Chiao, Y.A., Dai, D.-F., Drake, R., Levy, N., Koh, W.J., Emond, M.J., Kruse, S., Marcinek, D., et al. (2015). Respiratory chain protein turnover rates in mice are highly heterogeneous but strikingly conserved across tissues, ages, and treatments. FASEB J. 29, 3582–3592. 10.1096/fj.15-272666.

32. Krishna, S., Arrojo e Drigo, R., Capitanio, J.S., Ramachandra, R., Ellisman, M., and Hetzer, M.W. (2021). Identification of long-lived proteins in the mitochondria reveals increased stability of the electron transport chain. Dev. Cell 56, 2952–2965.e9. 10.1016/j.devcel.2021.10.008.

33. Bomba-Warczak, E., Edassery, S.L., Hark, T.J., and Savas, J.N. (2021). Long-lived mitochondrial cristae proteins in mouse heart and brain. Journal of Cell Biology 220. 10.1083/jcb.202005193.

34. Stiburek, L., Cesnekova, J., Kostkova, O., Fornuskova, D., Vinsova, K., Wenchich, L., Houstek, J., and Zeman, J. (2012). YME1L controls the accumulation of respiratory chain subunits and is required for apoptotic resistance, cristae morphogenesis, and cell proliferation. Mol. Biol. Cell 23, 1010–1023. 10.1091/mbc.E11-08-0674.

35. Kohler, A., Carlström, A., Nolte, H., Kohler, V., Jung, S.-J., Sridhara, S., Tatsuta, T., Berndtsson, J., Langer, T., and Ott, M. (2023). Early fate decision for mitochondrially encoded proteins by a molecular triage. Mol. Cell 83, 3470–3484.e8. 10.1016/j.molcel.2023.09.001.

36. Szczepanowska, K., Senft, K., Heidler, J., Herholz, M., Kukat, A., Höhne, M.N., Hofsetz, E., Becker, C., Kaspar, S., Giese, H., et al. (2020). A salvage pathway maintains highly functional respiratory complex I. Nat. Commun. 11, 1643. 10.1038/s41467-020-15467-7.

37. Hornig-Do, H.-T., Tatsuta, T., Buckermann, A., Bust, M., Kollberg, G., Rötig, A., Hellmich, M., Nijtmans, L., and Wiesner, R.J. (2012). Nonsense mutations in the COX1 subunit impair the stability of respiratory chain complexes rather than their assembly. EMBO J. 31, 1293–1307. 10.1038/emboj.2011.477.

38. Szilagyi, Z., Michon, P., Pardo-Hernández, C., Sãcultanu, M., Miralles Fusté, J., Griffin, A.M., Charifson, P.S., Phan, C., Shi, Y., Kern, G., et al. (2026). A selective inhibitor reveals roles for LONP1 in mitochondrial proteostasis and OXPHOS dependencies. Life Sci. Alliance 9, e202503535. 10.26508/lsa.202503535.

39. Pöverlein, M.C., Jussupow, A., Kim, H., and Kaila, V.R. (2026). Protein-induced membrane strain drives supercomplex formation. Elife 13. 10.7554/eLife.102104.

40. Hughes, C.S., Moggridge, S., Müller, T., Sorensen, P.H., Morin, G.B., and Krijgsveld, J. (2019). Single-pot, solid-phase-enhanced sample preparation for proteomics experiments. Nat. Protoc. 14, 68–85. 10.1038/s41596-018-0082-x.

41. Chambers, M.C., Maclean, B., Burke, R., Amodei, D., Ruderman, D.L., Neumann, S., Gatto, L., Fischer, B., Pratt, B., Egertson, J., et al. (2012). A cross-platform toolkit for mass spectrometry and proteomics. Nat. Biotechnol. 30, 918–920. 10.1038/nbt.2377.

42. Kong, A.T., Leprevost, F. V, Avtonomov, D.M., Mellacheruvu, D., and Nesvizhskii, A.I. (2017). MSFragger: ultrafast and comprehensive peptide identification in mass spectrometry-based proteomics. Nat. Methods 14, 513–520. 10.1038/nmeth.4256.

43. Demichev, V., Messner, C.B., Vernardis, S.I., Lilley, K.S., and Ralser, M. (2020). DIA-NN: neural networks and interference correction enable deep proteome coverage in high throughput. Nat. Methods 17, 41–44. 10.1038/s41592-019-0638-x.

44. Derks, J., Leduc, A., Wallmann, G., Huffman, R.G., Willetts, M., Khan, S., Specht, H., Ralser, M., Demichev, V., and Slavov, N. (2023). Increasing the throughput of sensitive proteomics by plexDIA. Nat. Biotechnol. 41, 50–59. 10.1038/s41587-022-01389-w.

45. Le Sueur, C., Rattray, M., and Savitski, M. (2024). GPMelt: A hierarchical Gaussian process framework to explore the dark meltome of thermal proteome profiling experiments. PLoS Comput. Biol. 20, e1011632. 10.1371/journal.pcbi.1011632.

46. Schwanhäusser, B., Busse, D., Li, N., Dittmar, G., Schuchhardt, J., Wolf, J., Chen, W., and Selbach, M. (2011). Global quantification of mammalian gene expression control. Nature 473, 337–342. 10.1038/nature10098.

47. Hammarén, H.M., Geissen, E.-M., Potel, C.M., Beck, M., and Savitski, M.M. (2022). Protein-Peptide Turnover Profiling reveals the order of PTM addition and removal during protein maturation. Nat. Commun. 13, 7431. 10.1038/s41467-022-35054-2.

48. Ritchie, M.E., Phipson, B., Wu, D., Hu, Y., Law, C.W., Shi, W., and Smyth, G.K. (2015). limma powers differential expression analyses for RNA-sequencing and microarray studies. Nucleic Acids Res. 43, e47. 10.1093/nar/gkv007.

49. Pérez Pañeda, L., Misic, J., Kadavá, T., Larsson, N.-G., and Heck, A.J.R. (2026). Investigating the relationship between ATP synthase and the TCA cycle by crosslinking mass spectrometry. Nat. Commun. 17. 10.1038/s41467-026-74730-5.

50. R Core Team (2024). A Language and Environment for Statistical Computing. Preprint at R Foundation for Statistical Computing.

